# Interkingdom interactions shape the fungal microbiome of mosquitoes

**DOI:** 10.1101/2023.08.11.552965

**Authors:** Shivanand Hegde, Kamil Khanipov, Emily A Hornett, Pornjarim Nilyanimit, Maria Pimenova, Miguel A. Saldaña, Charissa de Becker, Georgiy Golovko, Grant L. Hughes

## Abstract

The mosquito microbiome is an important modulator of vector competence and vectoral capacity. Unlike the extensively studied bacterial microbiome, fungal communities in the mosquito microbiome (mycobiome) remain largely unexplored. To work towards getting an improved understanding of the fungi associated with mosquitoes, we sequenced the mycobiome of three field-collected and laboratory-reared mosquito species (*Aedes albopictus, Aedes aegypti,* and *Culex quinquefasciatus*). Our analysis showed both environment and host species were contributing to the diversity of the fungal microbiome of mosquitoes. When comparing species, *Ae. albopictus* possessed a higher number of diverse fungal taxa than *Cx. quinquefasciatus,* while strikingly less than 1% of reads from *Ae. aegypti* samples were fungal. Fungal reads from *Ae. aegypti* were <1% even after inhibiting host amplification using a PNA blocker, indicating that this species lacked a significant fungal microbiome that was amplified using this sequencing approach. Using a mono-association mosquito infection model, we confirmed that mosquito-derived fungal isolates colonize and for *Aedes* mosquitoes, support growth and development at comparable rates to their bacterial counterparts. Strikingly, native bacterial taxa isolated from mosquitoes impeded the colonization of symbiotic fungi in *Ae. aegypti* suggesting interkingdom interactions shape fungal microbiome communities. Collectively, this study adds to our understanding of the fungal microbiome of different mosquito species, that these fungal microbes support growth and development, and highlights that microbial interactions underpin fungal colonization of these medically relevent species.

## Introduction

The microbiome profoundly influences many phenotypes in a host. In mosquitoes, much of the focus in this area has centered on how bacterial microbiota play an important role in mosquito biology, particular in relation to vector competence or how bacteria can be exploited for vector control [1–4]. Many of these studies have examined how the bacterial microbiome influences mosquito traits important for vectorial capacity, including growth, reproduction, and blood meal digestion [5–9]. While these studies provide convincing evidence that microbes can influence traits important for vectorial capacity of mosquitoes [9, 10], the role of the fungi on mosquito biology is understudied and less well understood.

Several studies have characterized the fungal microbiome in different mosquito species using culture-dependant and -independent methods [11–27]. In general, these studies indicate the majority of fungal taxa that colonize mosquitoes are within the *Ascomycota* and *Basidiomycota* phyla [16, 19, 22, 28–31]. Shotgun metagenomic sequencing of *Cx. pipiens*, *Culisetra incidens,* and *Olchelerotatus sierrensis* uncovered a diverse array of fungal taxa in mosquitoes, but only two fungal genera, *Cladosporium* and *Chromocliesta,* were present in multiple mosquitoes [13]. Amplicon sequencing of bacterial and fungal microbiomes of *Ae. aegypti* found fewer eukaryotic taxa compared to bacterial, although the majority of eukaryotic reads in mosquitoes were designated to gregarine parasites, rather than fungal species [18]. While our appreciation of the fungal community is expanding, we have a poor understanding of its functional relavance or interactions with other members of the microbiome.

Fungal community composition and abundance appear to be influenced by several factors, similar to their bacterial counterparts [28]. Aspects that appear to affect fungal microbiota include habitat, host species, diet, and pathogen infection [16, 22, 23, 30, 31]. For instance, in the tree hole mosquitoes *Ae. triseriatus* and *Ae. japonicus,* both blood feeding and La Cross virus infection were shown to reduce fungal richness [17]. Like the bacterial microbiome, mycobiome community structure varies between mosquito species and habitats [16–19, 27] and fungal diversity is seen between mosquito tissues [19, 22, 30]. While it is evident that mosquitoes possess diverse fungal taxa, sequence based assessment of the fungal microbome can be challenging due to inadvertent amplification of the host. To overcome these challenges, methods to selectively amplify the fungal sequences at the expense of host sequence have been accomplished [11].

Fungi can influence mosquito phenotypes that have important ramifications for vectorial capacity. For instance, the presence of a common mosquito-associated *Ascomycete* fungus *Penicillium chrysogenum* in the midgut of *An. gambiae* enhances the mosquito’s susceptibility to *Plasmodium* infection [30]. Similarly, *Talaromyces* fungus increased *Ae. aegypti* permissiveness to dengue virus infection [31], while *Beauverua bassiana* reduces vectorial capacity of *Ae. albopictus* to Zika virus [32]. Other studies have examined the effect of yeast on mosquito development and survival, which are traits that could influence vectorial capacity. Supplementation of *Saccharomyces cerevisiae* or native yeast strains supported the development of *Cx. pipiens* [22], although there was a strain-specific effect on the overall growth and development [12]. Recent advances in rearing approaches have enabled mono-association infections to be undertaken whereby a single (or group) of microbe(s) is inoculated in to germ-free L1 larvae to enable mosquito growth and development [7, 23, 33, 34]. While studies using mono-axenic rearing approaches have focused on the influence of the bacterial microbiome on their ontogeny [7, 23, 35–38], the ability of fungal isolates native to mosquito fungi have not been evaluated using this innovative mosquito rearing approach.

To address these gaps in our knowledge regarding fungal-host association in mosquitoes, we used high-throughput sequencing to examine the fungal microbiome of *Ae. aegypti, Ae. albopictus* and *Cx. quinquefasciatus* mosquitoes caught in the field or reared in the lab. Using gnotobiotic infection approaches, we reared these mosquitoes mono-axenically with fungal isolates to examine colonization and effects on mosquito development. Our results provide insights into the role of the environment on the composition and abundance of the fungal microbiome, microbe-microbe interactions in mosquitoes, and the influence of native fungal isolates on mosquito life history traits.

## Material and Methods

### Mosquito samples and high-throughput sequencing

We used the DNA from *Ae. aegypti*, *Ae. albopictus,* and *Cx. quinquefasciatus* samples either collected from the field or reared in the lab for high-throughput sequencing to examine the fungal microbiome [35]. The field collection of mosquitoes were followed as described previously[35]. To characterize the fungal microbiome of these mosquito species, the internal transcribed sequence (ITS) was sequenced. The region spanning ITS2 was sequenced according to the Illumina metagenomic sequencing protocol. Libraries were prepared following the amplicon protocol which includes the use of indexes from the Nextera XT Index Kit v2 (Illumina). Library preparation was done according to Illumina amplicon protocol (Illumina) (Table S1, ITS primers) [39]. Libraries were sequenced on the MiSeq System with the MiSeq Reagent Kit v3 (Illumina, Catalog No. MS-102-3003). All MiSeq runs were performed with a run configuration of 2 x 251 cycles for PNA blocker PCR samples (see next section) and 1×501 cycles for all other samples. To enable the calculation of error-rate metrics and to increase nucleotide base diversity for more accurate base-calling, all libraries were spiked with 5% PhiX Control v3 (Illumina, Catalog No. FC-110-3001). The NCBI Genbank accession number for the raw sequencing data reported here is PRJNA999749.

### PNA blocker PCR with microbiome samples

To block host amplification, PNA blocker was designed and synthesised (PNA Bio, USA). The PCR was performed with 1 µM of each primer (Table S1), 2 µM PNA,1X KAPA master mix (NEB) and 50 ng of template DNA. The PCR conditions were as follows: 3 min at 95°C for initial denaturation; 30 cycles of 30 sec at 95°C, 30 sec at 70°C, 30 sec at 55°C, 30 sec at 72°C, 5 min at 72°C, then 30 sec at 70°C clamping step for PNA. The product was digested with *Sph*I which cuts the fungal ITS amplicon but not the region in mosquitoes (Fig. S1). The PCR products were purified and sequenced as described above.

### Bioinformatic analysis

To identify the presence of known fungi, sequences were analyzed using the CLC Genomics Workbench 12.0.3 Microbial Genomics Module. Reads containing nucleotides below the quality threshold of 0.05 (using the modified Richard Mott algorithm) and those with two or more unknown nucleotides were excluded and finally the sequencing adapters were trimmed out. Reference based OTU picking was performed using the UNITE v7.2 Database [40]. Sequences present in more than one copy but not clustered in the database were placed into *de novo* OTUs (97% similarity) and aligned against the reference database with an 80% similarity threshold to assign the “closest” taxonomical name where possible. Chimeras were removed from the dataset if the absolute crossover cost was three using a kmer size of six. Additionally, OTU’s were reclassified using BLASTn 2.7.1+[41] against the nt nucleotide collection database. The blast results were used for taxonomic categorization of the origin of ITS sequences between those from the host, metazoan, and fungi. Alpha diversity was measured using Shannon entropy (OTU level), rarefaction sampling without replacement, and with 100,000 replicates at each point.

### Isolation and identification of fungal isolates from mosquitoes

Homogenates of five adult female mosquitoes were from *Ae. albopictus* (Galveston strain) and *Cx. quinquefasciatus* (Galveston strain) were plated on Brain Heart Infusion (BHI) agar (BD Difco), Yeast Peptone Dextrose (YPD) agar (BD Difco), malt extract agar (BD Difco), Yeast Malt agar (BD Difco), and Sabouraud Dextrose Broth (BD Difco). Colonies were purified by streaking a colony on a fresh agar plate and incubated at 30 °C for 2 days and transferred to 22-25°C until colonies to appeared before proceeding with culturing in the respective media. Five colonies from each growth media type were screened based on the colony characteristics (Table S2). Genomic DNA was isolated and PCR used to amplify ITS as the way to identify the isolated fungi. The PCR was completed using 1x reaction buffer (NEB), 200 µM dNTPs, 1 µM of each primer (Table S1), and 1U of Taq DNA polymerase (NEB). The PCR conditions were an initial denaturation of 1 min, 30 sec at 95°C, then 35 cycles of 30 sec 95°C, 30 sec at 55°C, 30 sec 72°C and a final extension of 5 min at 72°C. The PCR products were separated on agarose gels before Sanger sequencing with ITS3 and ITS4 primers. Sequences were analysed using the BLAST tn NCBI database.

### In vitro growth analysis of fungal isolates

The growth of *R. mucilaginosa*, *C. oleophila*, *S. cereviciae* and *L. thermotolerance* were undertaken by culturing in liquid YPD medium at 28°C. Overnight cultures of fungal isolates were diluted 1:100 in YPD medium and were grown at 28°C for 48 hours. The growth was assessed by recording OD at 600nm at 0, 2, 4, 8, 24 and 48 hours (Fig. S2). The assay was done in five replicates and repeated twice.

### Mosquito mono-association infection with fungi

Mono-association (MA) rearing was used to assess the colonization of fungi in absence of a natural microbiome. Axenic L1 larvae were generated as described previously [7, 35]. The 45 axenic larvae (N=15 per flask) were infected with 1×10^7 cfu/ml fungi *R. mucilogenosa, C. oleophila, L. thermotolerance, S. cerevisiase* and *C. neteri* bacteria. Fungi *R. mucilogenosa, C. oleophila, L. thermotolerance* are the culturable fungi present in the lab colonies of *Ae. albopictus* and *Cx. quinquefasciatus* mosquitoes and C. neteri is the abundant culturable bacteria found in the laboratory *Ae. aegypti* mosquito colony. All the procedures related to mono-association infection of mosquitoes were undertaken in a sterile environment and sterility was verified by plating larval water on LB agar plates. The mono-associated larvae were fed with sterile fish food at the concentration of 20 ugm/ml. The axenic L1 larvae without microbes have slow growth rates and do not reach pupation. For the mono-associated infections, larvae were maintained in the T75 flask till they reached pupae stage and the pupae were transferred to a container to eclose into adults. The adults were maintained on sterile 10% sucrose solution untill they were harvested for CFU quantification. The infection in the T75 flasks were maintained till day 16 by this time most of the larvae had pupated. To quantify their fungal or bacterial symbionts loads, we surface sterilized L4 larvae with 70% ethanol for 3 min and 2 times 1X PBS for 5 min. Larvae were then homogenized and plated on YPD agar for fungi and LB agar for bacteria. After incubation for 2 days, colonies were counted. Five larvae from each flask (total N=15) were tested for CFU analysis. Both bacterial and fungal quantification were done from the same larval and adult sample. Time to pupation and the percentage of L1 larvae to reach adult stage were recorded to determine the effect of fungi on mosquito growth and development. Time to pupation was recorded as the day when pupae were collected from the flask post infection. The number of adults emerged from each flasks (N = 15 larvae per flasks) were recorded and the percentage of L1 that emerged as adults was calculated. To assess the interkingdom interactions between native microbiome and fungi, *Ae. aegypti* mosquitoes were also infected with *R. mucilogenosa, C. oleophila, L. thermotolerance, S. cerevisiase* either in mono-association or in conventional rearing settings. The bacteria *C. neteri* was used as a control for inter-microbial interactions which we described in our previous study [35]. To assess the interkingdom interactions between fungi and bacterial microbiome, we did the fungi and bacteria infection of mosquito with and without native microbiome. All the procedures relating in in the interkingdom interactions study were followed as did for the mono-association infection.

### Fungal qPCR analysis

We used qPCR to determine the fungal load in *Ae. aegypti*, *Ae. albopictus,* and *Cx. quinquefasciatus* using 18S rRNA primers and probes [42]. PCRs consisted of 50-100 ng of DNA, 1µM of. each primer (Table S1), 225 nM of the TaqMan probe (Table S1) 1% formamide, 1X Platinum Quantitative PCR SuperMix-UDG w⁄ROX (Invitrogen Corp.) and molecular biology grade water. We used the following PCR conditions: 3 min at 50°C for UNG treatment, 10 min at 95°C for *Taq* activation, 15 sec at 95°C for denaturation, and 1 min at 65°C for annealing and extension for 40 cycles. We used host S7 or actin gene specific primers as endogenous control. The relative fungal copies were compared to host genome copies.

### Statistical analysis

All statistical analysis of the CFU and mosquito growth analysis data were done using GraphPad Prism software. We first D’Agostino & Pearson test and Shapiro-Wilk tests to assess the normal distribution of data. The data sets which followed Gaussian distribution were analysed by One-Way ANOVA. The nonparametric Tukey’s multiple comparison test was performed on data sets which did not follow Gaussian distribution. The prevalence data were analysed by a Fisher exact test with 2×2 matrix where number of infected and uninfected for each treatment was compared with every other treatment for each mosquito species. P-value 0.05 was considered significant.

## Results

### Fungal microbiome sequencing and analysis

We sequenced the ITS2 region from field-collected and lab-reared mosquitoes to characterize their fungal microbiome. Across all samples, we obtained 9,310,520 reads and recorded, on average, 155,175 reads per mosquito sample. However, similar to other high throughput sequencing (HTS) studies characterizing the fungal microbiota in eukaryotic hosts [11, 43, 44], our attempts were hampered due to the amplification of host or metazoan sequences. This was most pronounced for *Ae. aegypti* where about 99% of the reads were nonfungal derived (Fig. 1), while *Cx. quinquefasciatus* and *Ae. albopictus* had an average 21% and 8% fungal reads, respectively. To block nonselective amplification in *Ae. aegypti* samples, we employed a PCR clamping approach using a PNA blocking probe. While we saw evidence of suppression of host ITS amplification in PCR-based assays (Fig. S1) and a large reduction of host ITS reads (38% reduced to 0%), this did not result in a substantial increase in fungal reads (Fig. 1; a change from 0 to 1%). PNA blockers have been previously used to exclude *Anopheles* 18S rRNA reads when sequencing [11] but we saw little difference in the fungal reads, mainly due to an increase in amplification of metazoan sequences as a percentage of the overall reads in the PNA blocker treatment (Fig. 1). We speculated that these *Ae. aegypti* lacked significant fungal communities and therefore we saw non-specific amplication of host DNA in this sample as there was a lack of fungi ITS template to amplify. To further address this we completed qPCR to estimate total fungal density in lab-reared mosquitoes using universal fungal primers. Here we saw significantly reduced fungal loads in *Ae. aegypti* compared to the other two mosquito species (Fig. S3; ANOVA with Dunn’s multiple comparison test, P<0.0001). Given the evidence for reduced fungal loads in *Ae. aegypti*, our attention then focused on examining the fungal microbiome of *Cx. quinquefasciatus* and *Ae. albopictus* mosquitoes (Table S3). Despite the fungal reads comprising a relatively small proportion of the overall reads in *Cx. quinquefasciatus* and *Ae. albopictus*, rarefaction curve analysis indicated that our sampling depth was sufficient to observe the majority of fungal OTUs in the majority of indivudal mosquitoes (Fig. S4).

**Figure 1.**
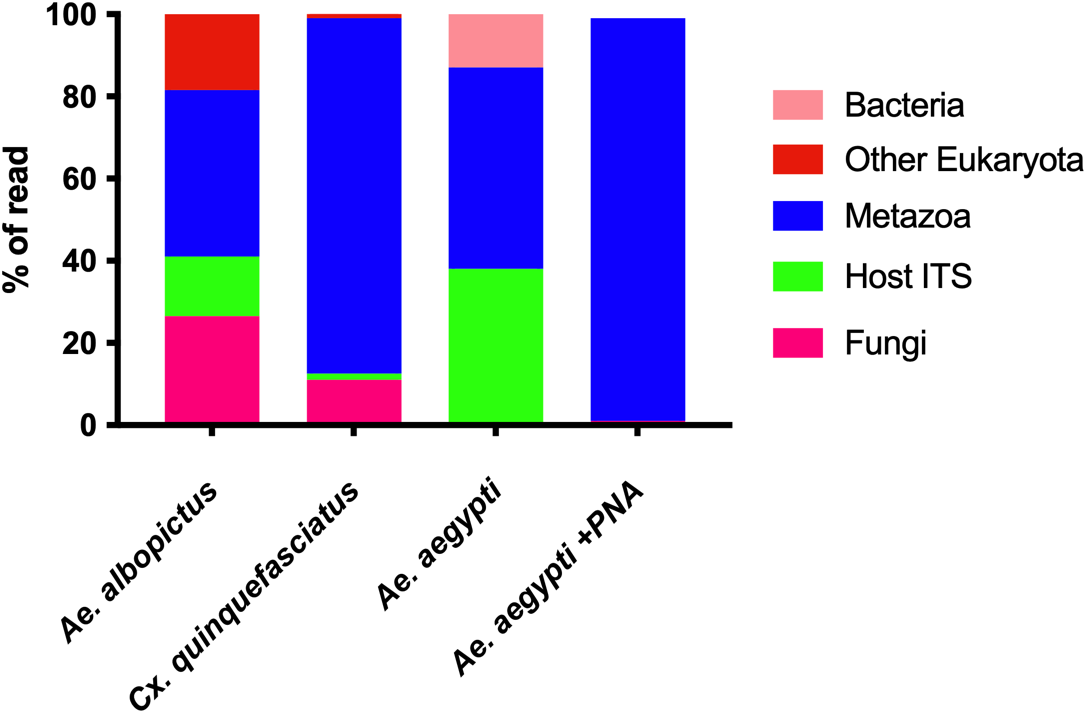
Average of percentage of reads from ITS2 sequencing. Average of percentage of ITS2 sequencing reads from *Ae*. *aegypti*, *Ae. albopictus* and *Cx. quinquefasciatus*. Reads were assigned to phyla. *Ae. aegypti* samples were sequencing again with the addition of a PNA blocker targeting the host ITS sequence. To generate average reads per species 11 laboratory reared and 22 field collected samples from *Ae. albopictus* and *Cx. quinquefasciatus* were analysed to generate average reads per species. For *Ae. aegypti* 22 field collected samples were assessed while 16 field collected samples were amplified with the PNA blocker.

### Fungal richness, diversity, and community structure

We examined the species richness of the fungal microbiome in *Cx. quinquefasciatus* and *Ae. albopictus* by evaluating the difference between field-collected mosquitoes caught in either the gravid (G) or BG traps. When comparing within each species, we saw no significant difference in the Shannon diversity between traps (BG or gravid traps, Fig. S5A; Tukey’s multiple comparison test, P > 0.05) for either species nor did we see significant differences between traps for beta diversity estimates (Fig. S5B and S5C; Bray-Curtis dissimilarity, P > 0.05). As such, we combined these mosquitoes for further analyses and considered them “field-collected”. When comparing between mosquito species, we found that the field-collected *Ae. albopictus* had significantly elevated Shannon diversity compared to *Cx. quinquefasciatus* (Fig. 2A; Tukey’s multiple comparison test, P < 0.05), but no difference was seen between species in lab-reared mosquitoes. Similarly, there was no significant difference in Shannon diversity when comparing within a species between environments (i.e. field vs lab; Fig 2A). This was also true for the number of OTUs with no difference within a species but *Ae. albopictus* had significantly more OTUs compared to *Cx. quinquefasciatus* regardless of environment (Fig 2B; Tukey’s multiple comparison test, P < 0.05). We then examined the community structure of the fungal microbiome using Bray-Curtis NMDS analysis. Overall, the fungal microbiome clustered distinctly with both species and environment were identified as significant factors (Fig. 2C; Bray-Curtis dissimilarity, P = 0.0009). This was predominantly driven by the field samples which, when analyzed separately, were significantly different between each species (Fig 2D; Bray-Curtis dissimilarity, P = 0.0009), but when mosquito species were reared in a common lab environment the fungal microbiomes were similar (Fig 2E; Bray-Curtis dissimilarity, P = 0.05295). When comparing field-caught and lab-reared mosquitoes, both *Ae. albopictus* (Fig 2F; Bray-Curtis dissimilarity, P = 0.002997) and *Cx. quinquesfaciatus* (Fig 2G; Bray-Curtis dissimilarity, P = 0.0009) had distinct microbiomes, indicating environmental factors contributing to the diversity of fungal communities.

**Figure 2.**
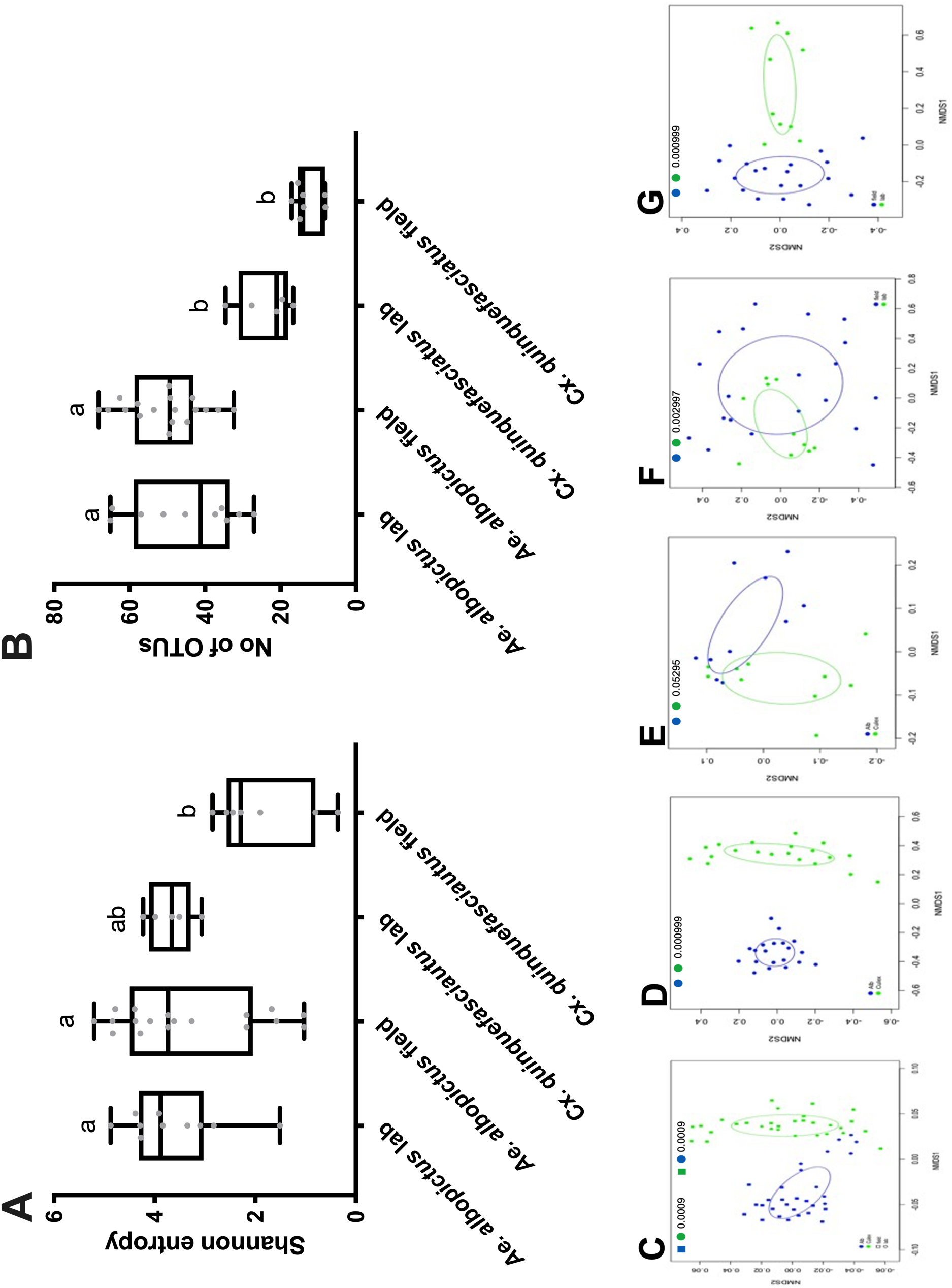
Alpha and Beta diversity analysis of fungal microbiome. Shannon entropy measuring abundance of fungal microbiome in *Ae. albopictus* and *Cx. Quinquefasciatus* (A). Number of operational taxonomic units represents species richness of fungal microbiome in *Ae. albopictus* and *Cx. quinquefasciatus* (B). Non-metric Multi-dimensional Scaling (NMDS) plots showing Bay-Curtis analysis of relative abundance of fungal OTUs. The fungal diversity was compared between laboratory and field samples between *Ae. albopictus* and Cx. *quinquefasciatus* (C). The fungal microbiome in the field collected samples (D) and laboratory reared mosquitoes (E) were compared between the two species. The fungal diversity between different environments (lab v field) was compared within *Cx. quinquefasciatus* (F) and *Ae. albopictus* (G). Numbers inside the graph indicates the p-value between groups. The field samples includes mosquitoes were collected in G and BG traps. Key for coloured squares and cicrles is within each panel. The data were analyzed by one-way ANOVA with Tukey’s multiple comparison test where P<0.05 considered significant.

Next, we examined the taxa present in each mosquito species. There were 244 fungal OTUs in mosquitoes, of which 76 and 97 were present above a 0.1% threshold in *Cx. quinquefasciatus* and *Ae. albopictus*, respectively (Table S3). While the majority of taxa were unidentified (Fig. 3 and S6), of the known OTUs, most were classified within the *Ascomycota* and *Basidiomycota* phyla, (Fig S6), which was similar to other studies [16, 18, 45]. *Saccharomycetaceae* were the most abundant in *Ae. albopictus* while the *Malasseziaceae* where dominant in *Cx. quinquefasciatus* (Fig 3A and S6). Unsurprisingly, considering the beta diversity analysis, the microbiomes of the lab-reared mosquitoes were comparable, however when examining the diversity between individuals, there was variation (Fig S6), which is also a feature of the bacterial microbiome [35]. In many cases, OTUs that were dominant in one individual were absent or at low abundances from others (Fig S5).

**Figure 3.**
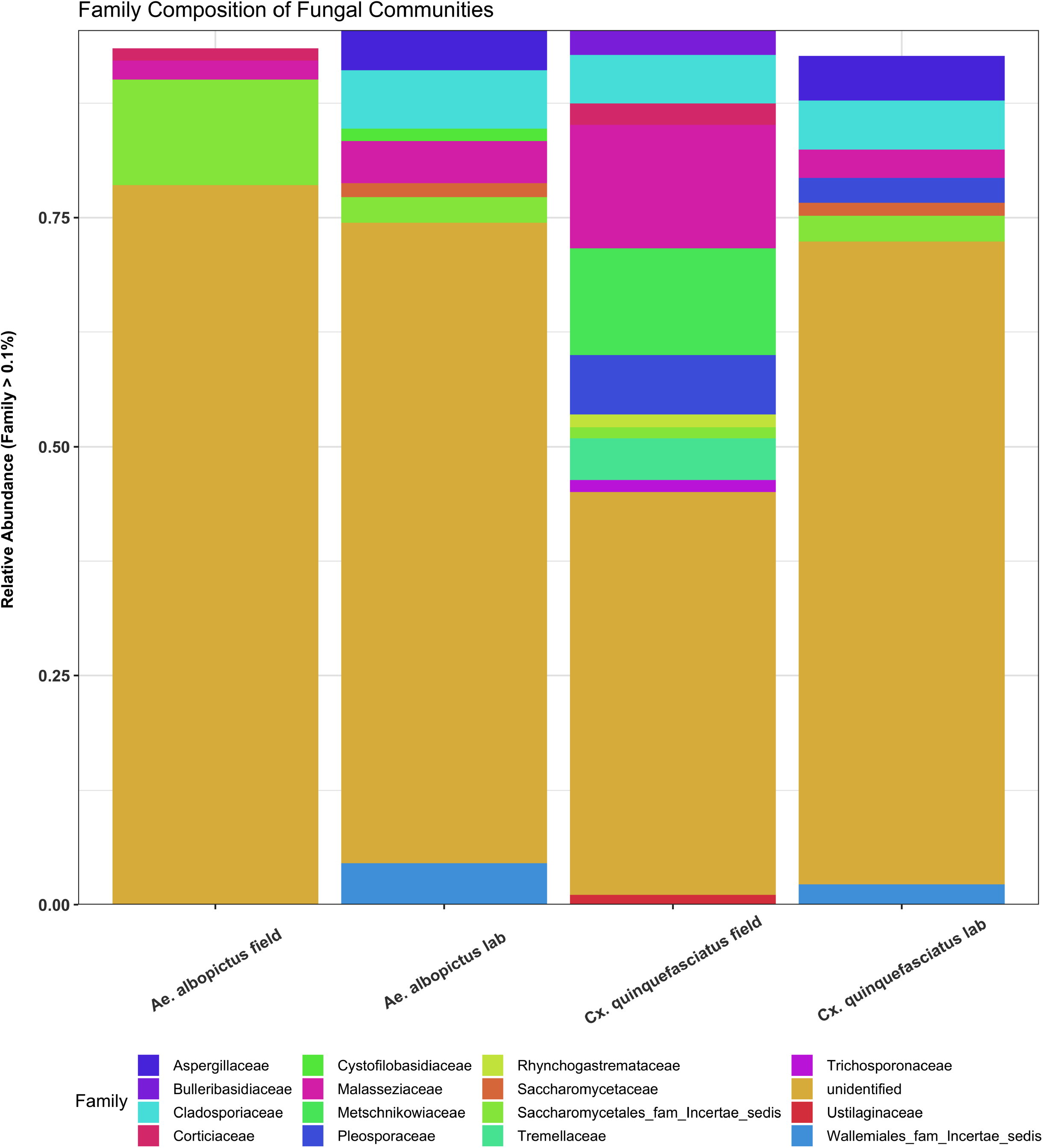
Relative abundance of fungal taxa. The relative abundance of fungal OTUs at family level with 0.01% cut-off between *Ae. albopictus* and *Cx. quinquefasciatus* field and laboratory samples.

### Fungal isolates colonize and supports mosquito growth in mono-association

Microbes are required for mosquito growth and development [7, 37]. Eukaryotic microbes such as the model yeast, *S. cerevisiae*, are known to promote larval growth [23], however it is not clear how symbiotic fungi affect mosquito growth and development. We cultured and identified symbiotic fungi from *Ae. albopictus* and *Cx. quinquefasciatus*. To determine if these native fungal taxa colonize mosquitoes and supported growth of their hosts, we reared mosquitoes in a mono-association using four fungal species. Three of these species, *Candida oleophila, Rhodotorula mucilagenosa*, and *Lachancea thermotolerans* were native mosquito isolates while the model yeast *Sachharomyces cerevisiae* was used as a positive control. The growth of mosquitoes infected with fungi was also compared to a native bacterial isolate, *Cedecae neteri*, which is a common bacterium present in our lab-reared *Ae. aegypti* and complements growth of mosquitoes in a mono-association [35]. When colonizing germ-free mosquitoes, fungi were more effective at colonizing *Ae. aegypti* and *Cx. quinquefasciatus* (Fig. 4A and 4C, circles, Fisher’s exact test, P>0.05) having high prevalance rates in adults while prevalance was reduced for all microbes in *Ae. albopictus* (Fig. 4B, circles, Fisher’s exact test, P<0.05). Intrigingly, colonization rates of 100% were observed in both larvae and adults of *Ae. aegypti* for all microbes (Fig. 4A, circles). Additionally, the native fungal densities were comparable to that of the symbiotic bacteria *C. neteri* (Fig. 4A, Dunn’s multiple comparition test, P<0.05). Both *C. oleophila* and *R. mucilaginosa* poorly infected adult *Ae. albopictus* despite infecting larvae (Fig. 4B, Dunn’s multiple comparition test, P<0.05). Similar to *Ae. aegypti*, the native fungal infection prevalence was 100% in larvae while there was no significant difference in the infection prevalence of microbes in adults (Fig. 4C, circles, Fisher’s exact test, P>0.05) although variable infection densities were observed in both life stages (Fig. 4C, Dunn’s multiple comparisonn test, P<0.05).

**Figure 4.**
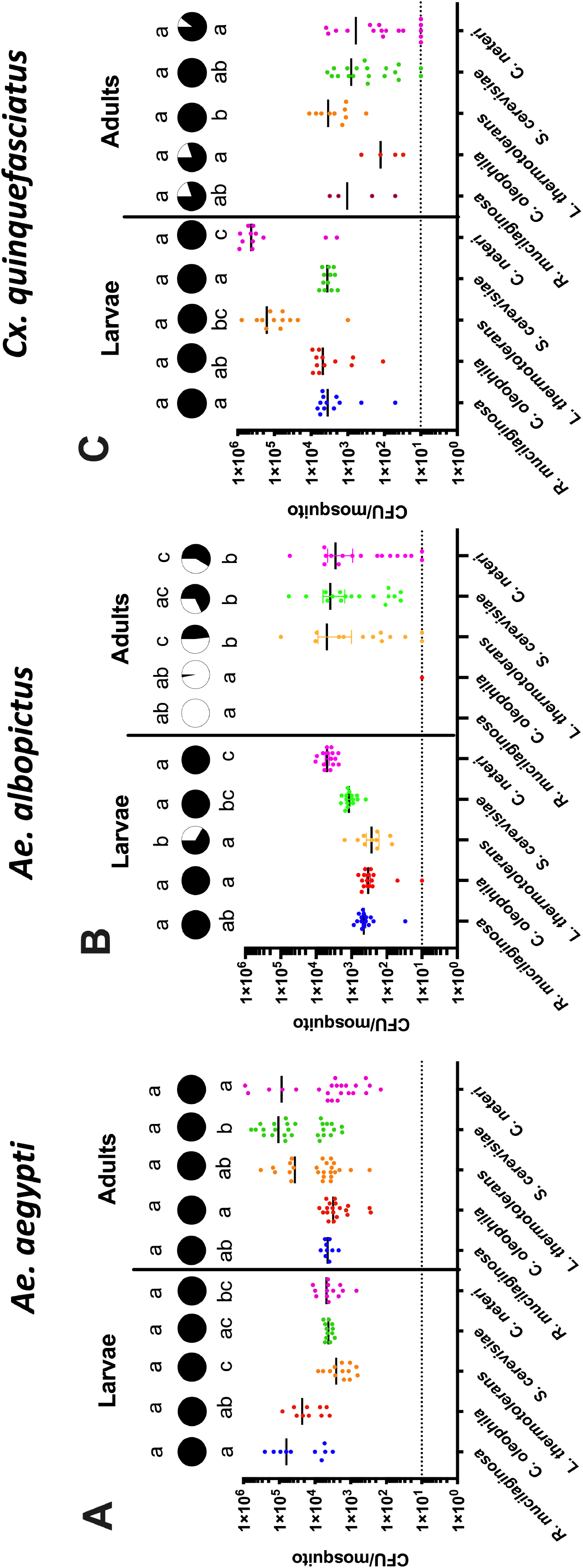
Fungal colonization of axenic mosquitoes. The scattered plot shows CFUs/mosquito of *Ae. aegypti* (A), *Cx. quinquefasciatus* (B) and *Ae. albopictus* (C) larvae and adults. The CFU data were analysed by Kruskal-Wallis Test with a Dunn’s multiple comparisons test. The circle above each scattered plot shows prevalence of infection for that treatment. Prevalence data were analysed by Fisher exact test. Letters above each scattered plot and prevalence circle indicate significance between the treatments. For all statistical analysis P<0.05 was considered significant. Sample size was N ≥ 10 for larvae and N ≥ 5 for adults – each dot on the graph represents an individual mosquito. The dotted horizontal line inidicates threshold detection limit.

### Mosquito development assay

Given bacterial microbiota can influence development we also determined the life history traits associated with mono-association infection. In *Ae. albopictus*, mosquitoes infected with *L. thermotolerans* had reduced times to pupation compared to the other native fungal microbes, while there was variability in pupation times in *Ae. aegypti* but no differences in *Cx. quinquefasciatus* between microbes (Fig 5A-C). We also measured the percentage of L1 larvae that reached adulthood in these mono-associations. In general, *Ae. albopictus* had higher rates of mosquitoes reaching adulthood for all microbes, while the percentage of *Culex* mosquitoes emerging as adults was below 40% for all fungal taxa (Fig 5D-F). In *Ae. aegypti* mosquitoes, *R. mucilaginosa* infections hadsignificantly different effects compared to the other two native fungi, while in *Ae. albopictus* its effects were only significantly different from *S. cerevisiae* (Fig 5D &E, Tukey’s multiple comparision test, P<0.05).

**Figure 5.**
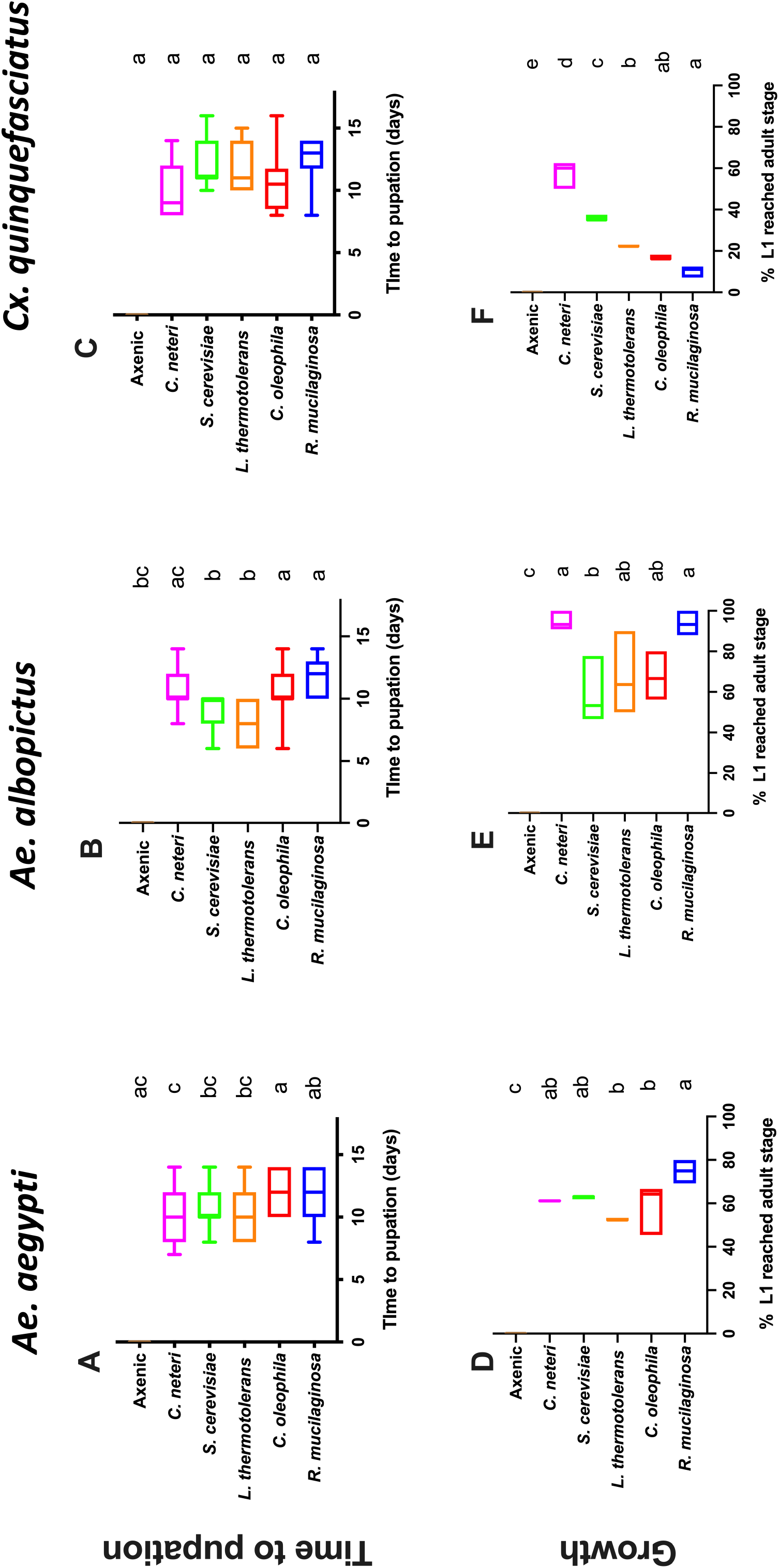
Life history traits in mono-association infections. Time to pupation of each species in mono-axenic associations (A-C). Data were analysed by one-way ANOVA with Dunn’s multiple comparision test. Growth was determined by percentage of L1 larvae to reach adulthood (D-E). Data were analysed by one-way ANOVA with Tukey’s multiple comparision test. None of the axenic larvae pupated and hence, the percentage to adulthood are zero for all axenic controls.

### Fungal infection in presence and absence of native bacterial microbiome

We have previously shown that colonization of symbiotic bacteria is influenced by members of the native bacterial microbiome [35, 46]. Given the ability of fungi to infected *Ae. aegypti* in a mono-association but the lack of fungal reads in field-collected mosquitoes, we speculated that bacteria may inhibit fungal infection. To determine if cross kingdom interactions influenced fungal colonization, we infected fungi into conventionally reared or axenic *Ae. aegypti*, which either possessed or lacked their native bacterial microbiome, respectively. Strikingly, we did not recover any fungal CFUs in either larvae or adults when the mosquitoes were grown conventionally in the presence of a native microbiome, however in stark comparison, fungal isolates were able to effectively colonize germ-free mosquitoes (Fig. 6, Mann Whitney Test, P<0.05). Intringuingly, the reduced colonization capacity of fungi of conventionally reared mosquitoes was seen in both larvae (Fig. 6A, Mann Whitney Test, P<0.05) and adults (Fig. 6B, Mann Whitney Test, P<0.05). In agreement with our previous study [38], the positive control, *C. neteri* also was more effective at colonizing germ-free mosquitoes compared to their conspecfic’s that possessed a conventional microbiome, however this effect here was more subtle compared to the almost complete blockage of fungi seen when mosquitoes had bacterial microbiota.

**Figure 6.**
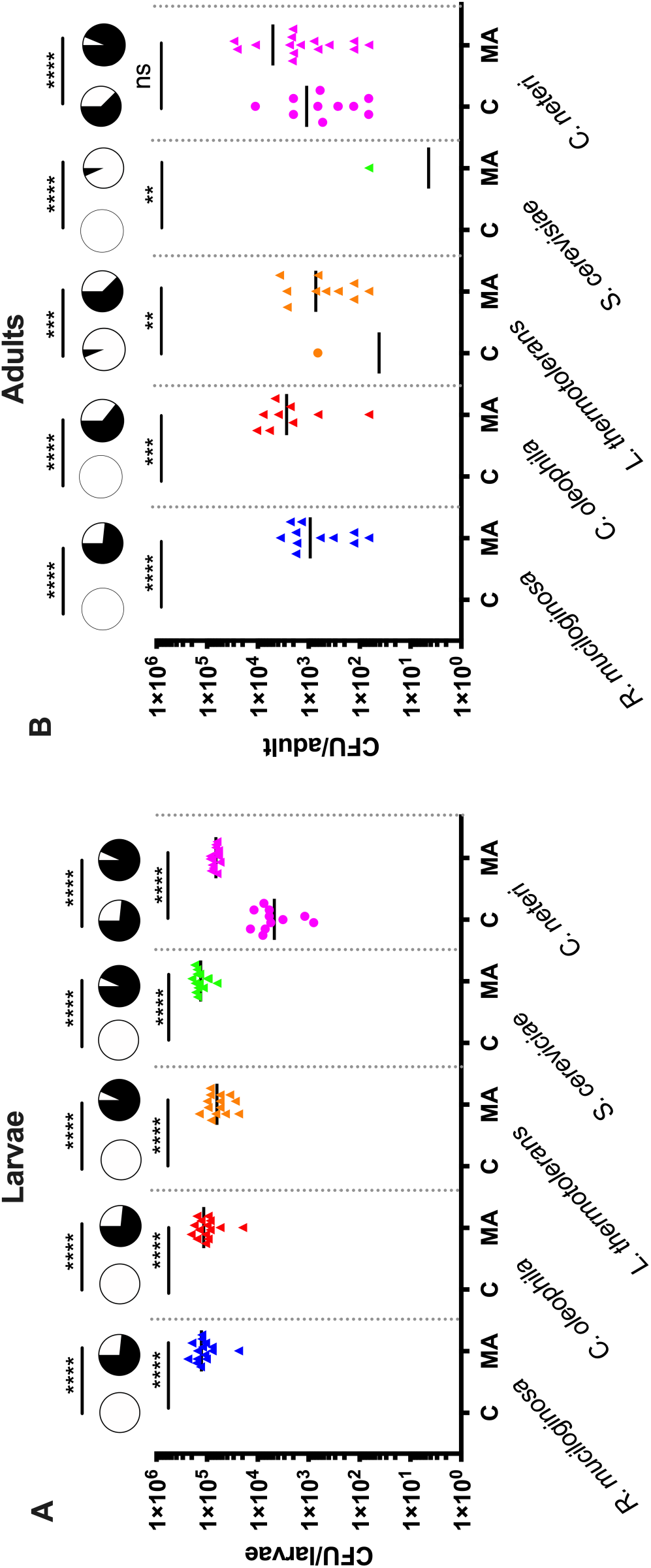
Fungal colonization in presence or absence of a native microbiome. *R. mucilaginosa*, *C. oliophila*, *L. thermotolerans* were incolulated into conventionally (C) reared *Ae. aegypti* mosquitoes that possessed their native microbiota or axenic germ-free mosquitoes to create a mono-association (MA). CFUs were quantified in **A)** L2-L3 larvae and **B)** three to four day old adults. The bacterium *C. neteri* was used as a positive control. A contamination control was undertaken by rearing axenic larvae without infection. These mosquitoes did not develop confirming sterility. The CFU/mosquito data were analysed by unpaired t test and prevalence data by a Fishers exact test. Asterisks (*) indicates significance, while ns denotes non-significant.

## Discussion

We characterised the fungal microbiome of *Ae. aegypti*, *Ae. albopictus and Cx. quinquesfaciatus* collected from different environments. Sufficient fungal reads were obtained from *Cx. quinquesfaciatus* and *Ae. albopictus* to evaluate their fungal microbiomes. In these species, we found the fungal composition varied substantially between species and environments. These findings were similar to other reports whereby environment has been shown to be a major determinant of fungal microbiome composition [16, 18, 19]. At the individual level, there was variability in the composition of fungal taxa within mosquitoes. Of the known taxa, *Malassezia*, *Saccharomcetales*, and to a lesser extent, *Candida* were fungi that were frequently seen in either species and other studies have identified these genera in mosquitoes suggesting they may commonly infect these vectors [13, 16, 29, 47, 48].

Strikingly, our sequencing data suggest that the fungal microbiome of *Ae. aegpyti* is dramatically reduced as we only observed a small fraction of fungal reads in these mosquitoes. Initially we speculated that the low number of fungal reads was due to preferential amplification of the host, and as such we used blocking PNA oligonucleotides to suppress host reads, in a similar fashion to other studies [11, 43, 44]. Despite our blocking primer reducing host ITS reads, there was no significant increase in the number of fungal reads, but rather an increase in off target host reads, indicating that these field caught mosquitoes lacked fungi at an amplifiable level. Supporting this finding, qPCR analysis of lab-reared *Ae. aegypti* found significantly reduced fungal densities compared to *Ae. albopictus and Cx. quinquesfaciatus*. Together these data indicate that these *Ae. aegypti* mosquitoes have a reduced fungal microbiome. Further studies are required to determine if this is consistent across other lab-reared or field collected *Ae. aegypti* mosquitoes.

Little is known about the capacity of members of the fungal microbiome to colonize their mosquito host. Although our sequencing data indicate *Ae. aegypti* lacked a robust fungal microbiome, specific taxa were able to colonize when infected into germ-free mosquitoes. The ability of germ-free mosquitoes to harbour fungi suggests that the reduced fungal load that we saw in *Ae. aegypti* by sequencing or qPCR was not due to an incompatibility between the fungal species and the mosquito, but rather due to microbial incompatibility. To empirically test this, we compared infection of fungal taxa in germ-free compared to conventially reared mosquitoes and found fungi infected the mosquitoes in absence of native microbiome. While the microbiome can be composed of a variety of microbes, we speculated that bacterial microbiota were interfering with fungal infections. We have previously identified several bacterial co-occurrence interactions in these mosquitoes and experimentally validated inter-bacterial interactions in co-infection studies [35, 46, 49]. However, fungal-bacterial co-occurance has not been exclusively investigated. Several other studies identified fungal and bacterial communities co-existing from individual mosquitoes, but these were not in *Ae. aegypti* [13–15]. More generally, the influence of bacteria-fungi interactions on colonization has been observed in diverse microbial systems including the soil microbiome, and the microbiota of livestock and humans [50–53], so further investigations of these interactions in mosquitoes are warranted.

Several studies have shown that the bacterial microbiome is required for mosquito growth and development [7, 38, 54]. Other eukaryotic microbes can also facilitate development including the model yeast *S. cerevisiae* and insect cells [23, 55]. Here we show that native fungal species that associate with mosquitoes also have the ability to support mosquito growth and development. We did observed developmental variation between fungal microbes and between mosquito species, however, *S. cerevisiae* had similar developmental rates compared to previous studies [23, 55]. Interestingly, we saw variability between replicates in terms of *S. cerevisiae* infections. These replicate experiements (Fig 4A [*S. cerevisiae* had high prevalance and density] and Fig 6 [lack of *S. cerevisiae* infection]) were performed on the same mosquito lines but reared at different institutions. Our most recent analysis of microbiome from these mosquito lines reared at these different insectaries revealed they possessed significantly different microbiomes [56] and given our findings regarding fungal-bacterial interactions, it is tempting to speculate that differences in the native microbiota were responsible for the variation in *S. cerevisiae* colonization. These findings will be important to confirm given that *S. cerevisiae* is being investigated for novel vector control strategies [57].

In summary, here we showed that *Ae. albopictus* and *Cx. quinquefasciatus* harbor fungal taxa as part of their microbiome, but, *Ae. aegypti* appear to lack mycobiome. The lack of fungal taxa in *Ae. aegypti* appears to be due to cross kingdom microbial interactions. Despite this, when the bacterial microbiome is removed, fungi can infected these mosquitoes and support their growth. Together, our findings have shed a light on an understudied aspect of the mosquito microbiome and shown that native fungal symbionts influence mosquito biology.

## Supplementary tables and figures legends

**Table S1.** Sequences of PCR primers used in the study

**Table S2.** Characteristics of mosquito derived fungal isolates. The size, color of the colony screened for each species isolated from the *Ae. albopictus* Galveston and *Cx. quinquefasciatus* colony. The fungal species were indentified by Sanger sequencing.

**Table S3.** Complete and filtered OTU table with relative abundance from each individual mosquito (*Ae. albopictus and Cx. quinquefasciatus*).

**Figure S1. PNA blocker PCR:** (Left) Schematic representation of PNA blocking PCR against host ITS. The amplicon was digested with *Sph*I, which specifically cuts fungal ITS. (Right) Agarose gel showing the PCR products done with *Ae. aegypti* laboratory samples in presence or absence of PNA blocker. The PCR product was digested with *Sph*I.

**Figure S2. In vitro growth analysis of fungi.** The fungal isolates were grown in YPD medium at 28 C for 48 hrs and OD_600_ was recorded at indicated time points. The experiment was repeated twice each with 5 replicates. The data were analysed by two-way ANOVA with Tukey’s multiple comparision test. The assay was done twice each in 5 replicates.

**Figure S3. Total fungal abundance.** The fungal load in the laboratory reared mosquitoes is analysed by qPCR using primer specific 18S rRNA gene and host endogenous gene S7 and Actin were used as control. The Ct values were normalized to host genes are represented in the graph. The data were analysed by one-way ANOVA with Dunn’s multiple comparision test. The P<0.05 considered significant.

**Figure S4. Rarefaction curve:** Alpha diversity species richness at intervals between 0 and 30,000 reads in each sample from different groups lab and field samples in *Ae. albopictus* and *Cx. Quinquefasciatus*.

**Figure S5. Abundance and diversity of fungal microbiome field samples. (**A)Alpha diversity analysis of fungal communities in *Ae. albopictus* and *Cx. quinquefasciatus* samples collected using gravid (G) and BG sentinel traps. The statistical significance was determined by one-way ANOVA with Tukey’s multiple comparison test. The P<0.05 considered significant. The diversity of communities in the G and BG samples of *Ae. albopictus* (B) and *Cx. quinquefasciatus* (C) were analysed by Bray-Curtis metric.

**Figure S6: Beta diversity analysis:**The detailed view of the comparison of abundance at family level between *Ae. albopictus* and *Cx. quinquefasciatus* field and laboratory samples.

## Acknowledgements.

We would like to thank the UTMB insectary core for providing the lab mosquitoes. GLH was supported by the BBSRC (BB/T001240/1, BB/V011278/1, BB/X018024/1, and BB/W018446/1), the UKRI (20197 and 85336), the EPSRC (V043811/1), a Royal Society Wolfson Fellowship (RSWF\R1\180013), the NIHR (NIHR2000907), and the Bill and Melinda Gates Foundation (INV-048598). SH was supported by a Director’s Catalyst Fund at the Liverpool School of Tropical Medicine and Royal Society Research Grant (RGS\R1\231156).

## Data accessibility

The NCBI accession number for the raw sequencing data reported here is PRJNA999749.

## Author Contributions

SH, KK, EAH, and GLH designed the experiments. SH, PN, MAS, and MP completed the experiments. KK, GG, SH, EAH and GLH undertook analysis. SH, KK, and GLH wrote the first draft and SH, KK, CDB, and GLH, edited the manuscript. All authors agreed to the final version. GLH acquired funding and supervised the work.

